# ProCAST: A Bioinformatics Suite for Mass Spectrometry-Based Protein Corona Proteomics Analysis

**DOI:** 10.64898/2026.05.08.723620

**Authors:** Huyju Mun, Maxwell Leamy, Anika Kaushik, Christopher A. Kieslich, Simone A. Douglas-Green

## Abstract

When nanoparticles are exposed to biological fluids, they spontaneously adsorb proteins, forming a protein corona that defines their biological identity and dictates cellular uptake, biodistribution, and toxicity. Characterizing protein coronas includes using proteomics approaches (e.g., LC-MS/MS) to identify proteins and generate vast lists of adsorbed proteins, often visualized via complex heatmaps. While heatmaps display data they do not offer heuristic guide, leaving the driving mechanisms of adsorption unknown. Moreover, interpretation of protein corona proteomics data remains limited by fragmented workflows, inconsistent preprocessing, and visual outputs that are often descriptive rather than readily interpretable. These conventional methods identify adsorbed proteins but fail to explain why specific proteins are selected or how they influence the particle’s biological fate. Here, we developed ProCAST (Protein Corona Analysis and Statistical Tool), an R-based framework for protein corona proteomics that integrates proteomics data, nanoparticle metadata, protein annotations, and multi-level visualization within a single analytical workflow. ProCAST facilitates abundant protein clustering based on sample conditions, sequence descriptors, property or protein correlations, and gene ontology-based functional visualization. It also distinguishes abundant proteins from frequent proteins, providing distinct layers of information from the same dataset. ProCAST was used to re-analyze previously published PAMAM G4 dendrimer-FBS datasets, demonstrating that ProCAST reproduces descriptor-level visualizations and offers new insights through clearer comparisons of functional patterns and hypothesis generation from dominant corona proteins. By organizing results as complementary views of the same dataset, ProCAST facilitates the shift of protein corona analysis from descriptive outputs toward structured, comparative, and experimentally testable interpretations.

## Introduction

Nanoparticles can be used as drug delivery systems to decrease toxicity, improve biodistribution and pharmacokinetics, offer controlled release, and enhance stability and solubility of therapeutic agents^1^. However, to ensure the safety and efficacy of nanoparticles, it is imperative to understand their interactions with biological systems and fluids. The dynamic interactions between biological interfaces and nanomaterials are a challenge and an opportunity for innovations in next-generation biomaterial development. According to the FDA^2^, nanomaterials can develop new or altered physicochemical properties in a biological context, which may significantly affect their safety and therapeutic efficacy. These changes in nanomaterial properties are driven by interactions with plasma proteins, leading to the formation of a protein corona that alters their targeting and transport. When nanoparticles are immersed in biological fluids, proteins^3^, lipids^4, 5^, metabolites^6^and carbohydrates^7, 8^ adsorb to their surfaces, forming a nanoparticle-protein complex called a protein corona. Protein coronas can alter the size, shape, and surface properties of nanoparticles, potentially affecting the nanoparticle’s intended targeting and therapeutic capabilities^8-13^. Limited understanding of how nanoparticles perform *in vivo*, especially due to bio-nano interactions, including protein corona formation^14-16^, could explain difficulties in successful clinical translation for nanomedicine applications^17-19^. Thus, a predictive program that models protein corona formation during the nanoparticle design process would eliminate these uncertainties and enable consideration of protein corona formation to optimize nanoparticle drug delivery. To achieve this level of prediction and optimization, a precise characterization of the protein corona is a necessary first step, as machine learning or artificial intelligence models highly depend on the quality of a training dataset.

Protein corona studies are typically designed around three steps: incubation, isolation, and analysis (**Figure 1A**)^20^. First, nanoparticles are incubated in the selected biological fluid, where incubation times range from 30 minutes to 1 hour at 37 °C, under static or lightly shaking conditions^20-24^. Next is isolation, where nanoparticle-bound proteins are separated from unbound proteins. Finally, isolated nanoparticle-bound proteins are characterized for protein concentration, changes in nanoparticle size or charge (due to bound proteins), and protein identity. Currently, there is no standard procedure to analyze protein-nanoparticle interactions; as such, researchers often rely on combining complementary characterization methods to overcome the limitations of individual techniques^25, 26^. Of the available techniques, mass spectroscopy (MS) remains the most prominent for characterizing the protein corona due to its ability to provide detailed compositional information^27^. However, protein corona characterization studies often show significant variability across research groups; this is in part due to how raw MS data is processed^28^.

**Figure 1.**
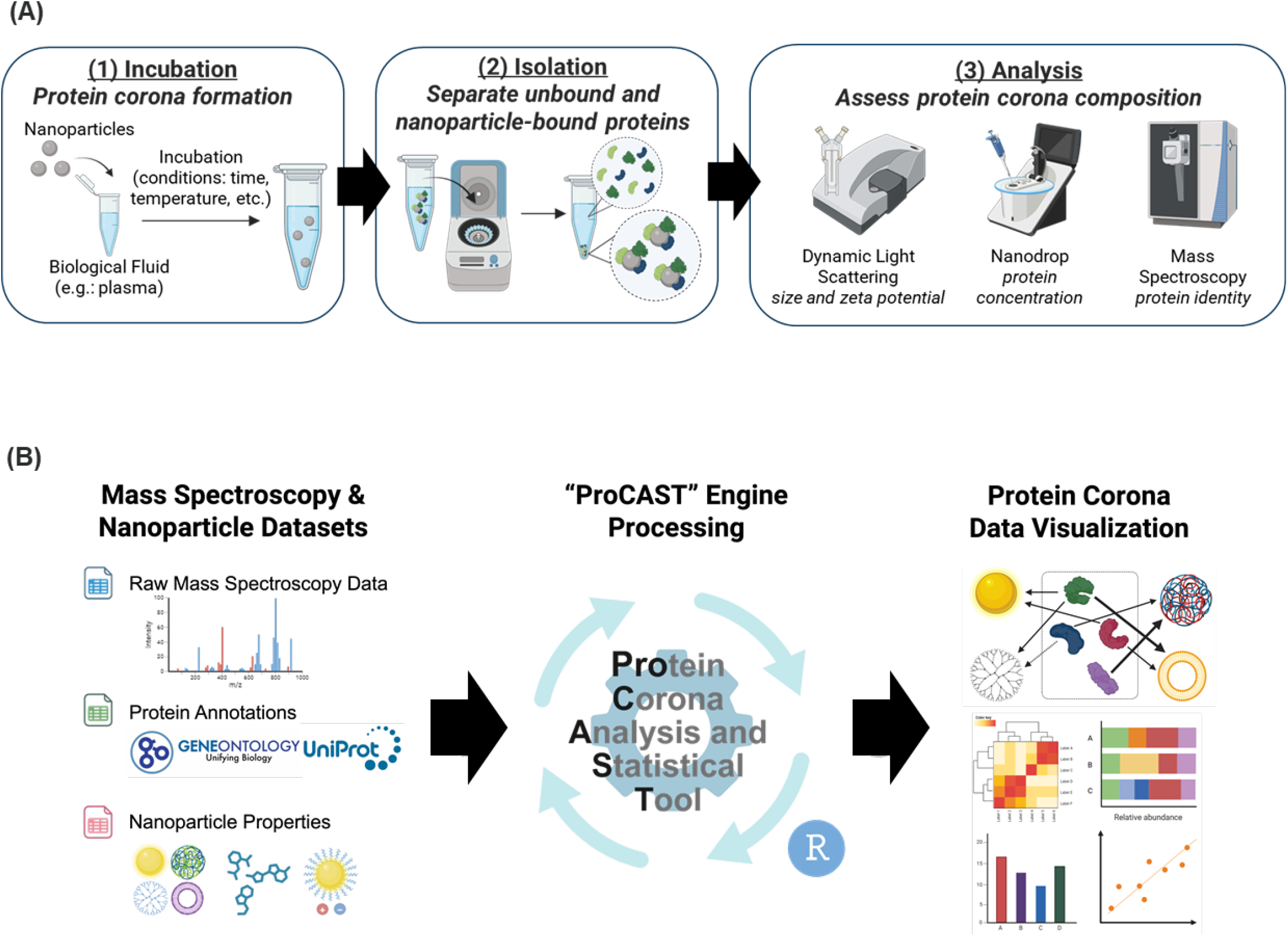
Overview of ProCAST for streamlined meta-analysis of protein corona proteomics data. **(A)** Typical workflow for protein corona characterization experiment: incubation, isolation, and analysis. **(B)** Proteomic data and nanoparticle/experiment metadata (e.g., size and charge) are input into ProCAST, which normalizes MS data, retrieves protein and gene ontology information, and calculates experiment and sequence-level descriptors, and generates resulting data visualizations including clustering and functional pathway analysis.

Protein corona proteomic analysis remains hindered by a lack of standardized practices for data processing and analysis^28^. Proteomic characterization relies on MS^29^, but approaches vary in how data are filtered, interpreted, and reported^30^. Inconsistencies limit cross-study comparisons, which slows progress in the development of nanomedicines. There have been recommendations for the bio-nano research community to use a “reporting standard”, MIRIBEL (Minimum Information Reporting in Bio-Nano Experimental Literature)^31^, which includes reporting material properties, biological properties, and experimental protocol details to improve the quality of research. This includes documenting the details of statistical and data analysis, with methods described in detail and any code used for analysis accessible through open-access tools.

Beyond the difficulties of standardizing raw MS data processing, the biological interpretation of proteomics data presents another major hurdle. Conventional protein corona analyses often remain descriptive^21-23,32^ even when substantial proteomics data are available. Ranked protein lists and pathway summaries can show what is present, but they do not always make it clear why certain proteins are represented, why others become dominant in specific conditions, how these shifts/patterns relate to nanoparticle properties, or how we can use this information for the next step. As a result, visual outputs are not always biologically or mechanistically interpretable at the level needed for rational follow-up. This creates a gap between protein identification and protein corona reasoning, where data can be generated but remain difficult to read in a way that supports comparison, hypothesis generation, and experimental design.

There is a range of bioinformatic tools to process proteomics data and elucidate the biological importance of protein composition. However, despite the availability of tools for protein identification, quantification, and annotation, there is no unified software platform that is tailored for protein corona proteome analysis. Consequently, researchers are compelled to rely on a complex patchwork that involves numerous software programs and protein and biological databases, such as UniProt, Gene Ontology (GO), and KEGG, to perform statistical analyses and data visualization on their MS proteomics data^20, 33, 34^. Researchers commonly employ a range of software, including ProtPram to obtain protein properties such as molecular weight and isoelectric point, Venny for Venn Diagram visualization, heatmapper for heatmap visualization, and platforms (e.g., DAVID, Cytoscape) to classify and visualize protein networks by biological process^35-38^. While some tools offer visualization functions for proteomic data, none combine physicochemical properties of nanoparticles and adsorbed proteins along with MS data in a single platform that conducts comparative analysis focused on protein coronas. A tool of this nature would be a major asset to researchers in the field of nanoparticle-based drug delivery, streamlining protein corona analyses.

To this end, we developed ProCAST (Protein Corona Analysis and Statistical Tool), a platform for bioinformatic data analysis of protein corona proteomes that enables statistical analysis and data visualization to characterize nanoparticle-protein interactions that best suit their research needs and experimental questions. ProCAST uses R and contributed packages to leverage augmented data matrices containing MS data, nanoparticle properties, and protein information to enable a user-friendly application to support standardized, reliable, and reproducible bioinformatics analysis of protein corona data generated by MS. ProCAST is capable of performing a meta-analysis using the same dataset utilized in other papers, it offers a more specific and comprehensive analysis, taking into account not only nanoparticles characteristics but also annotations of adsorbed proteins and providing a more intuitive and hypothesis-generating visualizations.

## Methods

### ProCAST Design and Workflows and Implemented Packages

ProCAST was developed by combining commonly known R-based data science, statistical, and bioinformatics packages, which enabled the integration of annotation, normalization, clustering, and visualizations into a unified framework tailored to protein corona proteomic analysis. Once loading sample and MS datasets, ProCAST employs the ‘UniProt.ws’ package to get protein annotations, including protein names, sequences, and physicochemical properties, connecting proteomics data to downstream descriptors and GO analyses. To overcome the challenges of missing data during preprocessing and to maximize annotation accuracy, ProCAST adopts a dual-database integration strategy. It prioritizes high-confidence GO annotations from UniProt (manually curated via SwissProt through literature review and experimental validation), which provides proteomic-centric information. This is then merged with data from Ensembl BioMart, accessed via the `biomaRt` package, which provides rapidly updated, genomic-centric information encompassing automated annotations, computational predictions, and manual curation. The `GO.db` package is utilized to access standardized GO term definitions, ontology categories (Biological Process, Cellular Component, and Molecular Function), and hierarchical structures. This complementary approach combining these resources enables ProCAST to minimize vulnerability to single-database failures; if an annotation is missing in one database, it can be retrieved from the other, ensuring that protein annotations remain up-to-date while providing a more comprehensive, consistent, and reliable interpretation of functional enrichment results across analyses. Using the ‘Peptides’ package, physicochemical properties such as molecular weight, charge, instability index, isoelectric point, amino acids composition were calculated based on sequence information obtained from UniProt, and these calculated values were restored in the ProCAST table. Protein abundance measurements (e.g., total spectral counts) were first normalized within each sample to generate sample-wise percent abundance relative to the total signal in that sample, after which frequent and abundant proteins were defined. This percent-normalized matrix was further used to rank proteins, summarize GO abundance, and visualize sample-condition-level clustering. Z-score normalization was additionally employed for cross-sample contrast in descriptor-based clustering and selected comparative visualizations, particularly where relative scaling provides a more intuitive grasp and discernible comparisons of the differential trends.

### Input Data Structure Requirements

ProCAST requires two structured input tables: a protein abundance table and a sample data table. The protein abundance table contains protein identifiers with quantitative values (e.g., total spectral counts, percent of total spectral counts) for each sample, while the sample data table maps each sample to experimental descriptors, including nanoparticle formulations, biological fluid, and experimental conditions, and may also include nanoparticle physicochemical properties such as size, charge, surface chemistry, or other reported variables. Sample identifiers were matched across the two input tables prior to downstream analysis. Previously published PAMAM dendrimer protein corona MS data were converted into ProCAST-compatible input tables for re-analysis by a separate data processing code written in R (Supplementary Information), while the sample data table was manually compiled.

### Protein Annotation Retrieval and Sequence-Based Description Calculation

Protein sequence, physicochemical annotation, and gene ontology information were obtained via accession-based mapping to external databases (e.g., UniProt, Ensembl, Gene Ontology). For the previously published dendrimer dataset, 10 protein entries could not be mapped directly because they were listed as either uncharacterized proteins or had been deleted from UniProtKB or Ensembl. In these cases, the original MS tables in the Supporting Information of the source study were manually reviewed, and the most reasonable replacement accession numbers were assigned using reported protein names, molecular weights, and related annotation fields. These replacements were implemented for cases where direct database mapping failed to match within the database. Details of the curated substitutions would be found in Supplementary Information. Manual harmonization of the accession number is the most critical step since sequence descriptors, GO annotation, and downstream grouping all rely on currently valid UniProt accession numbers. Any alteration in these identifiers would compromise the underlying biological and experimental interpretation.

### Normalization and Frequent and Abundant Protein Definitions

The purpose of normalization is to minimize technical noise among samples and emphasizing true biological significance. While trimmed mean normalization (trimmed mean of M-values; TMM) is commonly employed in RNA-seq to reduce the influence of extreme outliers by eliminating the top and bottom 10% of data, we found it less suitable for our proteomics dataset for two reasons. First, MS data consist of continuous intensity measurements while RNA-seq data consist of discrete gene read counts, and therefore, can only take on integer values. Second, the protein corona is a complex biological system where trace-level proteins, pathways despite their low relative abundance, might play a pivotal role in triggering cellular responses or mediating certain biological. Consequently, applying TMM could lead to the removal of the lowest 100% of the data by classifying it as “technical noise”, thereby risking the loss of essential “rare proteins” that determine the biological behavior of nanoparticles. As a result, total sum normalization was adopted, as it offers the most intuitive approach for preserving the relative abundance of all detected proteins. Data normalization was carried out in two stages: sample-wise percent normalization, which divides total spectral counts of individual protein by the total signal within each sample to account for concentration variations, and Z-score normalization, which standardizes the data by centering it around the mean and scaling by the standard deviation to facilitate direct comparison of differential trends across samples. Frequent proteins were defined as proteins with their highest cumulative abundance across all samples, whereas abundant proteins were defined as proteins with the highest maximum percent abundance observed in any sample after sample-wise percent normalization.

### Clustering Analyses

ProCAST includes four hierarchical clustering views: sample-condition-level clustering, sequence descriptor-based clustering, protein-abundance-based clustering, and functional-based clustering to preserve a different layer of structure in the dataset. Because protein corona matrices are often sparse, cosine distance was used rather than standard distance metrics (e.g., Euclidean distance), which can yield misleading results when datasets contain many zero values, while Euclidean distance was used for treating sequence descriptors data. Sample-level clustering used the percent-normalized matrix to compare overall corona similarity across nanoparticle and biological fluid conditions. Sequence descriptor-based clustering used a protein-by-descriptor matrix composed of sequence-derived physicochemical properties and amino acid composition features. Correlation-based clustering of abundant proteins used protein–protein abundance relationships across samples. Pairwise abundance correlations were converted to a correlation-derived distance-like matrix and visualized by hierarchical clustering to identify co-varying protein groups. Together, these clustering views preserve condition-level structure, descriptor-level structure, and coordinated abundance structure as distinct but related outputs.

## Results and Discussion

### Overview of ProCAST Design and Analytical Workflow

ProCAST was designed as a tool to characterize bio-nano interactions, with an emphasis on streamlining and standardizing the protein corona proteome analysis (**Figure 1B**). This allows the user to choose from an array of functionalities to identify biologically relevant information and generate plots and figures summarizing their findings. As such, the initial workflow and program functionalities were developed using the types of analyses and visualizations commonly used in protein corona characterization. Data visualization of protein corona composition was selected based on what is commonly reported; this includes reporting clustering heatmap of protein abundance and gene ontology^20, 22, 33, 39, 40^, top hit proteins^20, 22^ and sorting proteins based on molecular weight^20^, theoretical isoelectric point^20^, and biological function^41^. We also use ProCAST to introduce advanced statistical methods for protein corona composition, such as principal component analysis and hierarchical clustering, to explore correlations between nanoparticle properties (i.e., size, surface charge, surface chemistry, and material) and protein identities.

Details about coding and R packages used to develop ProCAST are included in the methods section, and **Figure 2** includes a detailed overview of the bioinformatics analysis workflow. To demonstrate the functionalities of ProCAST, we used previously published data where studies characterized the protein corona on polyamidoamine (PAMAM) dendrimers^22^. Briefly, dendrimers with varying degrees of PEGylation (using PEG MW 500kDa) were incubated in FBS (10% and 100%) for 1 hour at 37 °C and run on native PAGE. Dendrimers and proteins were identified in two bands on native PAGE, a top band of unbound free dendrimer and a “shielded band” containing dendrimers whose positive charge was shielded by PEGylation and protein-nanoparticle interactions. These bands were then cut and digested, and the resulting peptides were analyzed by MS to identify proteins. The purpose of analyzing this dataset was not only to reproduce published summaries but also to examine whether the same dataset could be reorganized into outputs that make data visualization and interpretation more explicit.

**Figure 2.**
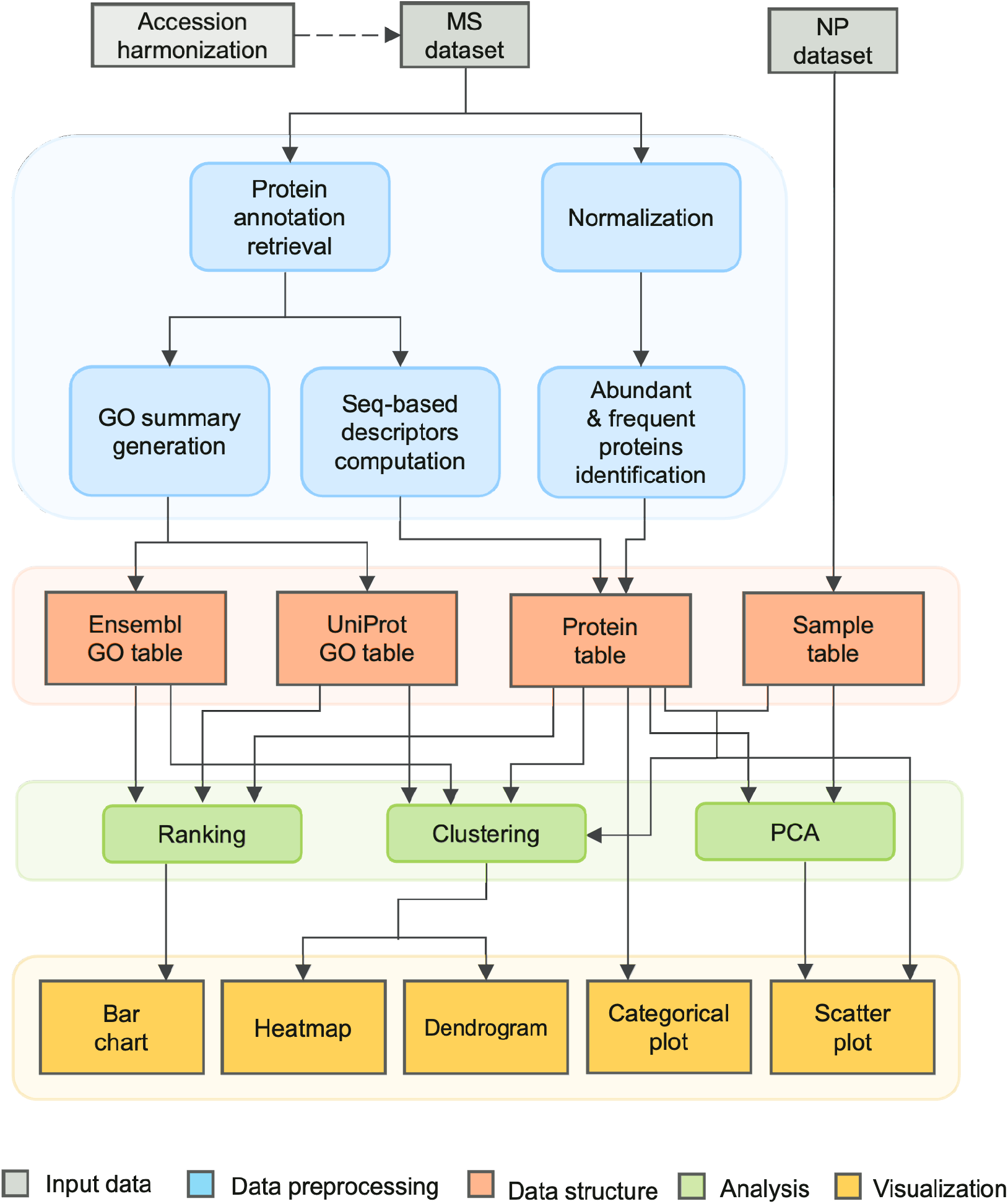
Detailed bioinformatics analysis workflow of ProCAST.

### Re-analysis of Published PAMAM Dendrimer Data to Demonstrate Reproducible Visualization

Using ProCAST, we reproduced plots quantifying the percentage of total spectrum counts using molecular weight and isoelectric point (**Figure 3**). ProCAST was also used to generate new graphs representing the Top 5 proteins for each condition (**Figure 4**); a feature which has been analyzed and reported in other protein corona analysis work^20, 42^. In the original workflow, these outputs required manual data organization and figure-by-figure preparation. In contrast, ProCAST generated the same class of descriptor-level views directly from the uploaded dataset, within a single consistent plotting framework which offers a streamlined, unbiased workflow for the user. Functional enrichment data using GO-based descriptors for molecular function, cellular component, and biological process were produced by ProCAST (**Figure 5**). The previously published dendrimer-protein dataset only includes biological process with data visualization to show global differences across dendrimers. With ProCAST, data is quickly processed for visualizing differences in proteins within a single dendrimer formulation. Analyzing data across groups provides a broad but limited summary for comparing condition-specific signatures across samples. However, ProCAST reorganizes the same information for a single nanoparticle, making it easier to analyze sample-specific functional patterns across nanoparticles.

**Figure 3.**
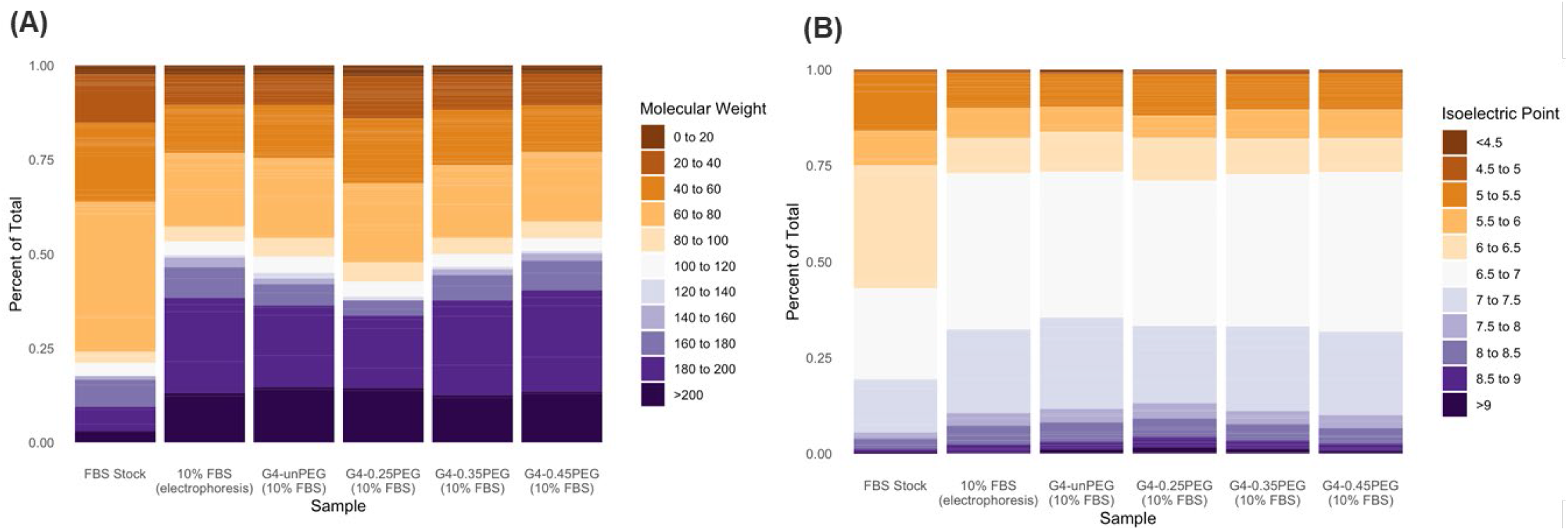
ProCAST provides standardized re-visualization of descriptor-based corona summaries from previously published PAMAM dendrimer datasets incubated in 10% FBS^22^. Sequence-descriptor-based data visualization for **A)** molecular weight and **B)** isoelectric point.

**Figure 4.**
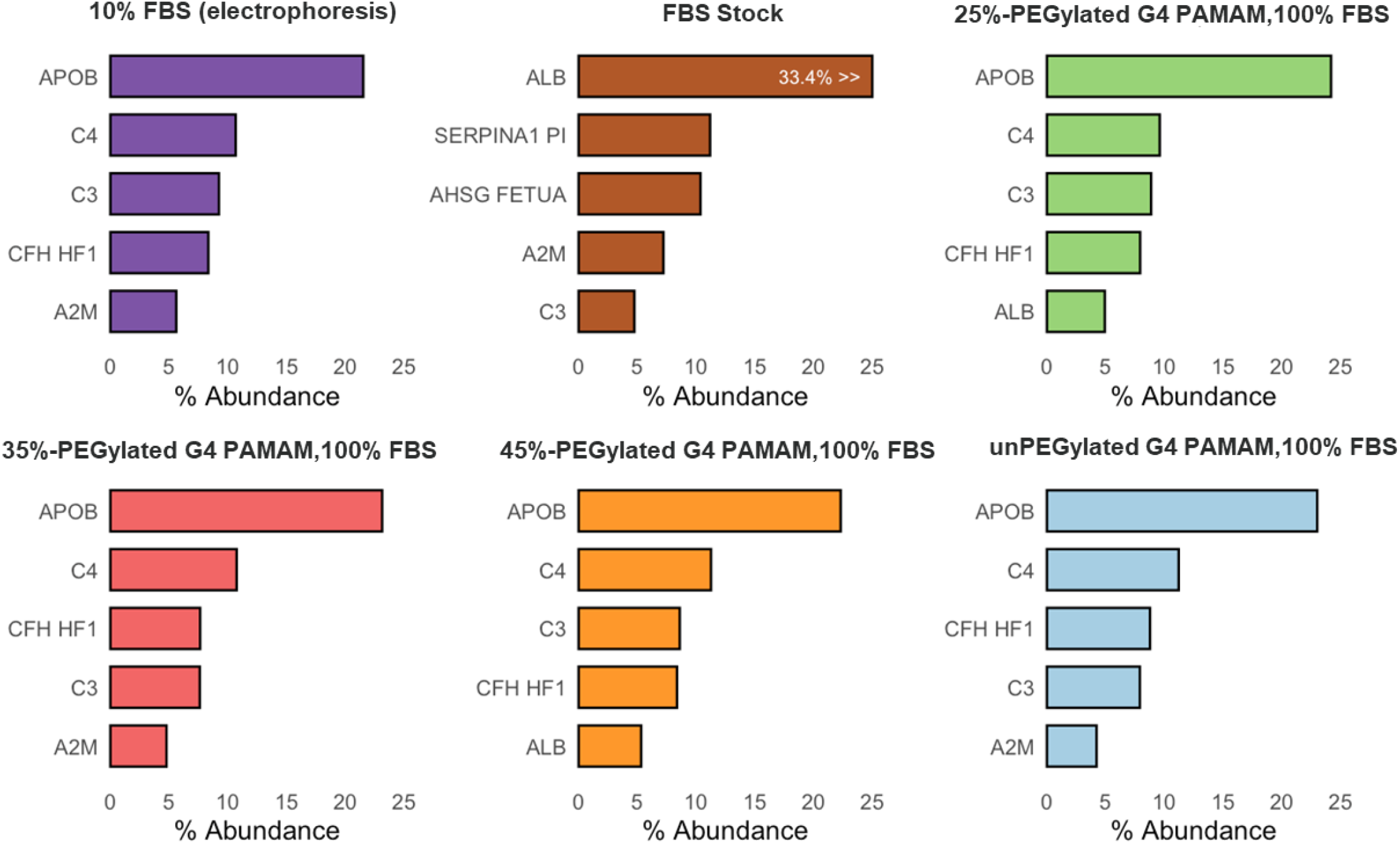
Abundance-based data visualization of the Top 5 proteins per dendrimer formulation using previously published dendrimer-FBS corona dataset incubated in 100% FBS ^22^. ProCAST enables intuitive comparison of top-ranked functional enrichment patterns of corona proteins

**Figure 5.**
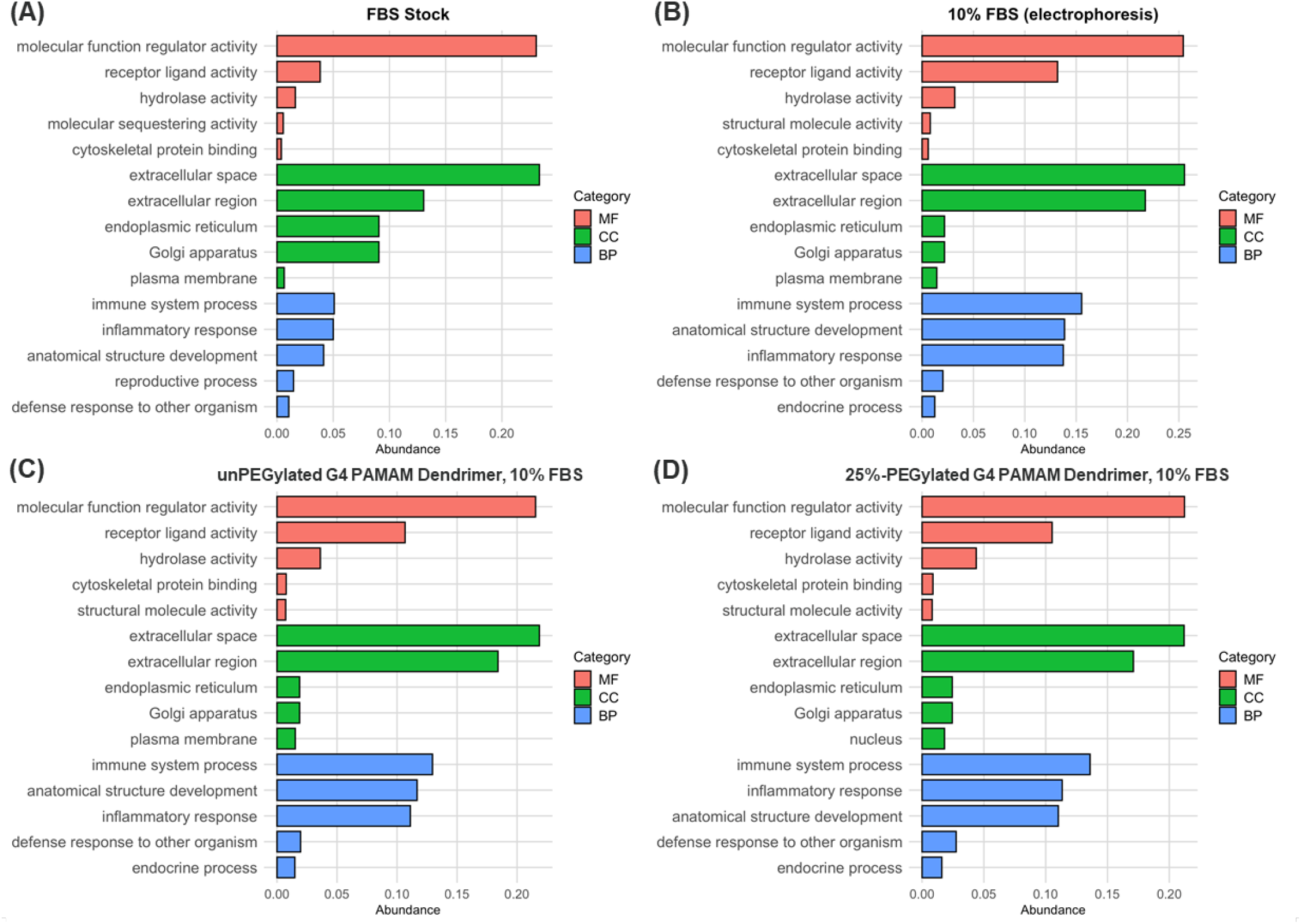
Functional enrichment data using GO-based descriptors for molecular function, cellular component, and biological process generated by ProCAST. Top 5 proteins by percent abundance for each sample of **previously published dendrimer-FBS corona dataset incubated in 10% FBS** ^22^: **A)** FBS stock, **B)** 10% FBS (electrophoresis), and **C) u**nPEGylated and **D)** 25%-PEGylated G4 PAMAM dendrimers.

### ProCAST Supports Multi-Level Interpretation of Protein Corona Composition

A main feature of ProCAST is the separation of frequent proteins from abundant proteins (**Figure 6A**). This distinction was intentional because recurrence across samples and dominance within a sample are not the same signal. Abundant proteins are those that reach peak quantitative dominance in one or more samples, while frequent proteins show high cumulative representation across the entire dataset. Treating these two metrics as identical would oversimplify the data, losing the potential distinct biological details each category provides. Here, an agnostic approach is used to report protein abundance and frequency which avoids assuming which metric is more informative. These metrics ensure a comprehensive and unbiased interpretation of the protein corona. ProCAST also includes multiple clustering views to preserve different kinds of structures in the same dataset. Sample-condition-level clustering (**Figure 6B**) compares overall corona similarity across samples representing different conditions for nanoparticle formulations and biological fluids. This provides a global view of how corona composition groups across experimental contexts. Sequence descriptor-based clustering (**Figure 6C**) groups proteins by intrinsic physicochemical features, allowing the dataset to be read at the level of protein classes rather than individual identities. Correlation-based clustering (**Figure 6D**) of abundant proteins provided additional insight by identifying co-varying protein groups across samples. These results identify organized patterns of protein-protein association rather than providing direct proof of a biological mechanism. Ultimately, ProCAST combines these perspectives to interpret the protein corona from multiple angles, including insights at the condition, property, and behavioral levels, serving as an instrumental framework for developing hypotheses rather than a definitive conclusion.

**Figure 6.**
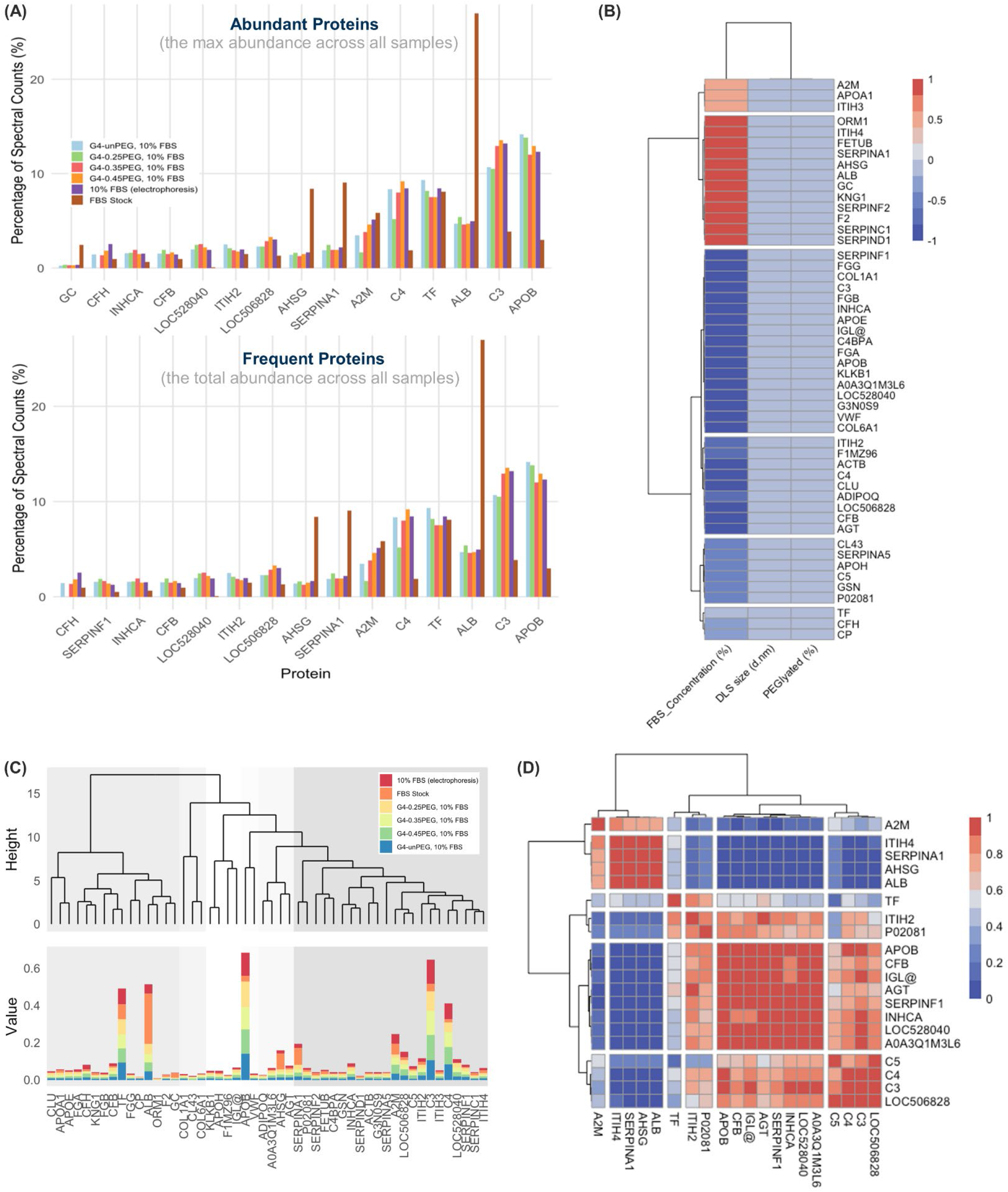
ProCAST enables multi-level interpretation of protein corona composition. **A)** Distinguishing the Top 15 abundant and frequent proteins across samples. **B)** Sample-condition-level clustering and **C)** sequence-descriptor-based clustering of abundant proteins. **D)** Correlation-based clustering of protein abundance, which shows co-varying abundance patterns among top proteins.

*ProCAST Incorporates GO-Based Visualization for Global and Sample-Specific Functional Interpretation* Protein functional interpretation was further extended through two complementary GO-based visualization modes, providing a deeper biological context. ProCAST summarizes GO-associated abundance patterns across all nanoparticle conditions (**Figure 7A**), making it easier to compare broad trends in molecular function, cellular component, and biological process. Additionally, ProCAST focuses on a single nanoparticle condition and highlights the dominant GO-associated signals within that sample (**Figure 7B**). Together, these two views enabled the simultaneous view of the shared structure of the dataset and the local structure of an individual corona profile. The advantage here is not just visual clarity, but interpretive clarity. This dual approach makes it easier to distinguish between shared trends and condition-specific functional signals. It allows researchers to interpret the protein corona at both the universal and local levels, capturing the full complexity of functional shifts across a variety of experimental settings.

**Figure 7.**
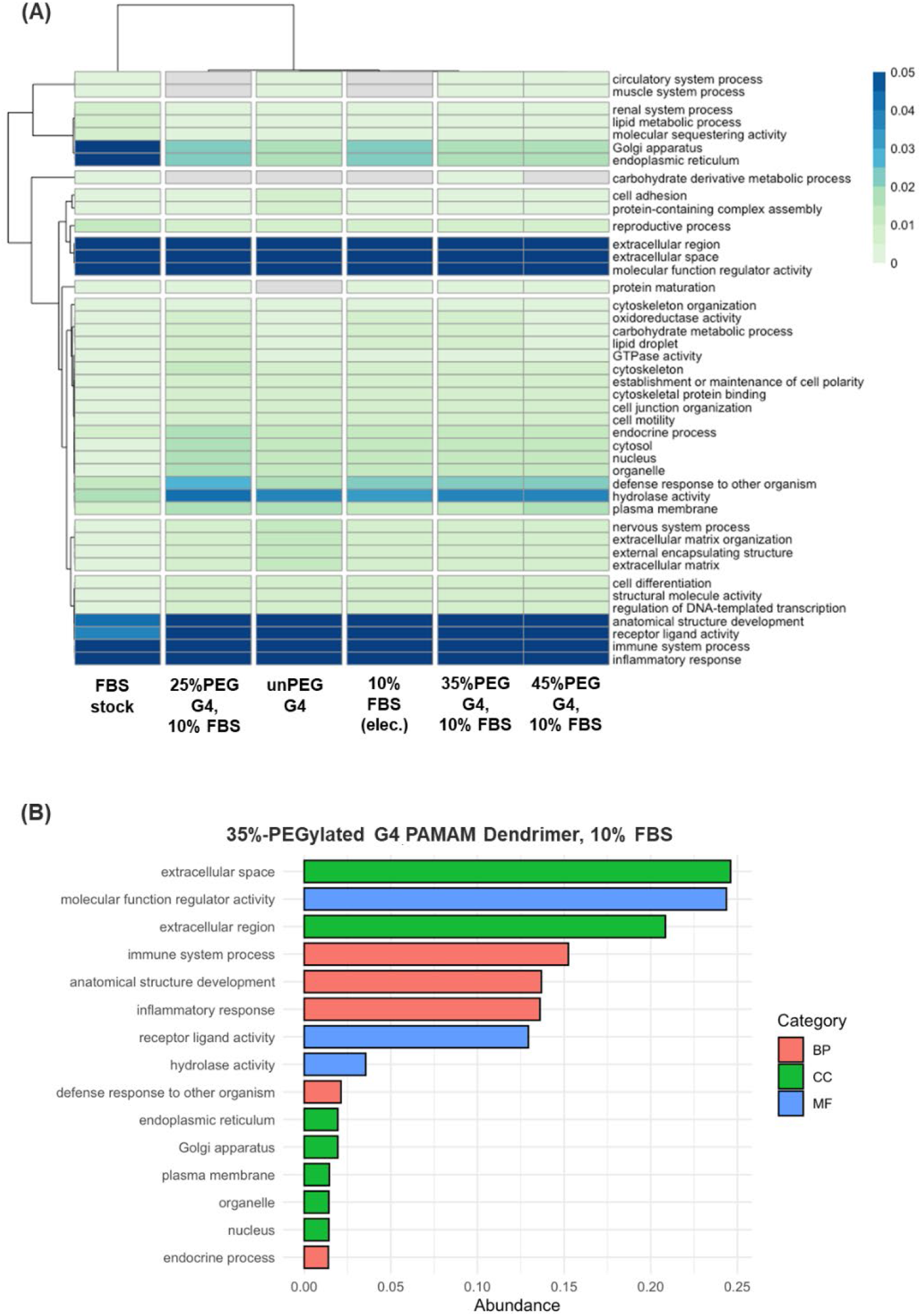
GO-based visualization supports global and sample-specific biological functional annotation. **A)** GO-based abundance patterns across nanoparticle conditions. **B)** Top GO-based signatures for a single nanoparticle condition.

### ProCAST as a Tool for In-depth Analysis and Hypothesis Generation

Data visualizations generated by ProCAST help researchers more thoroughly analyze protein corona data for improved functional and biological interpretation. Data visualization for abundant vs frequent proteins (**Figure 6A**) and sequence-based descriptor clustering (**Figure 6C**) showed similarity in abundance of proteins like Apolipoprotein B (APOB), Complement C3, C4, and transferrin (TF). APOB and C3 have been previously identified as abundant in coronas of nanoparticles with primary amines as surface functional groups^35, 43^. Complement proteins C4 and C3 are known inflammatory response proteins^44, 45^. The previously published analysis provides a useful overview, however this analysis is broad and limited for comparing condition-specific signatures across samples.

ProCAST reanalyzed the same proteomic data into condition-resolved GO-based visualizations (**Figures 5 and 7**), making it easier to systematically compare shared trends and sample-specific functional patterns across nanoparticle conditions. For example, in this dataset, unPEGylated and 25% PEGylated dendrimers share common biological patterns where protein coronas contain extracellular, immune-related, and signaling-associated terms. Through supporting system-level functional analysis, ProCAST moves beyond identifying which proteins adsorb to nanoparticle surfaces to elucidating biological mechanisms that inform how protein coronas affect immune response and downstream biological interactions. This feature is important because limitations in protein corona analysis arise not from a lack of data generation, but insufficient comparison of complex datasets. Conventional analysis approaches identify what proteins are adsorbed but, does not contextualize functional implications. ProCAST enables researchers to connect protein corona composition to biological insights regarding nanoparticle fate and response.

Outputs from ProCAST can be more useful when interpreted as hypothesis-generating rather than hypothesis-confirming. For example, when analyzing the top protein rankings in **Figures 4, 6A, and 6C** it was observed that albumin was the dominant protein in the FBS stock and remained among the top corona-associated proteins in several PEGylated dendrimer conditions, but not in the unPEGylated dendrimer condition. Although this finding does not establish improved biocompatibility, it highlights a formulation-dependent signal worth further testing. Beyond data visualization, ProCAST facilitates hypothesis generation by revealing formulation differences in protein corona composition that suggest testable biological differences across nanoparticles. By structuring and organizing data for direct comparison, ProCAST also helps clarify what experimental conditions need to be tested in future studies.

Taken together, these results show that ProCAST does more than develop figures in a more efficient format. By combining standardized visualization with comparative and condition-aware outputs, the platform enhances the interpretability of previously published protein corona datasets. In this way, ProCAST may support both retrospective re-analysis of literature datasets and forward-looking hypothesis generation for nanoparticle design.

### Limitations and Implications of ProCAST and Potential Research Suggestions

While ProCAST offers significant advantages for protein corona proteome analysis, there are limitations to consider. First, correlation- or clustering-based associations among proteins do not establish whether proteins co-adsorb directly onto the nanoparticle surface or whether sequential protein-on-protein adsorption events contribute to the observed patterns. Also, ProCAST outputs remain dependent on the quality, and completeness of the input dataset, which means results can be varied depending on the data users provide, as is the case with other machine learning and artificial intelligence tools. We initially attempted to incorporate fold change plotting, which is commonly reported in protein corona analysis studies, into ProCAST. However, we chose not to include this feature in the first iteration of ProCAST to prevent misleading interpretations or distorted information that could arise from flawed experimental data in the FBS stock baseline used for fold change calculations. While reviewing the literature, studies reported a change in corona composition, which may reflect not only nanoparticle physicochemical properties but also time-dependent alterations in the biological fluid itself, including proteolytic activity^46^ or other compositional remodeling during incubation^47-49^. For this reason, to improve interpretation, experimental conditions should include initial biological fluid controls and, when available, time-matched incubated biological fluid controls. In addition, although the field urgently needs standardized experimental protocols for protein corona studies, complete harmonization across laboratories remains challenging because differences in sample preparation, instrumentation, and MS preprocessing can yield different analytical outcomes even under the same samples. Lastly, while ProCAST supports more interpretable exploration of corona datasets, it does not by itself validate predictive relationships or establish mechanistic causality between nanoparticle properties and adsorption outcomes. These questions will require future experimental validation, benchmarking across various samples and biological fluids datasets, and potentially integration with predictive modeling approaches as a larger protein corona library is built.

Researchers have devoted efforts to establish databases to report proteomes (e.g., ProteomeXchange^50^) and protein corona specific databases (e.g., Protein Corona Database (PC-DB)^39^ and PROTCROWN^51^). However, these repositories primarily focus on the submission and dissemination of protein corona data and systematic analysis pipeline that enables researchers to immediately analyze the data is not yet sufficiently established. In 2025, Condchola, et al. reorganized and integrated 83 studies containing various nanoparticle types and biological fluids from 2000 to 2024 to analyze the global trends of protein corona meta-dataset ^39^. This study demonstrated that nanoparticle size, material type, and surface charge are the factors exerting the most critical influence on corona formation. Furthermore, by employing machine learning models such as LightGBM and XGBoost, the researchers predicted macroscopic patterns such as trends in dominant protein changes and GO enrichment. These prior studies have made contributions toward offering a broad overview of the universal trends within the field of protein corona research, what might be termed the “forest”. However, there is still the unresolved challenge of a lack of resolution regarding “zoomed-in” protein corona interpretations focused on specific sample sets that allow one to closely examine the individual “trees.”

Overall, ProCAST reduces manual analytical variability and provides a more consistent platform for systematic, user-friendly analysis of protein corona proteomics data all within a single workflow. With a single upload of nanoparticle and proteomic meta-dataset, it automatically retrieves physicochemical properties and gene ontology information from various databases commonly used in the proteomics research field, distinguishes frequent and abundant proteins. Additionally, ProCAST generates exploratory visualizations, including three different clustering approaches (sample condition-based, sequence-based, and abundance protein-based clustering), physicochemical property-dependent protein corona composition, and comparative universal and local trends of gene ontology. Validation of ProCAST using previously published datasets demonstrated that even the same dataset can yield different insights depending on how it is analyzed and visualized. This highlighted that ProCAST goes beyond conventional descriptive proteomic analysis, which is limited to identifying what proteins are present, and provides hypothesis-generating capability by enabling users to infer why certain proteins are present and how this information can guide the next experimental step.

While ProCAST was initially designed for protein corona proteome analysis, it is intended to support the entire bio-nano interfaces community. This universal platform is not restricted to nanoparticles and can be utilized in characterization studies in other emerging areas of research on organoids, extracellular vesicles, and implants. Since ProCAST requires only MS and sample datasets, it provides an accessible framework for researchers who wish to perform more thorough bioinformatics analysis but lack substantial coding or bioinformatics expertise. Thus, ProCAST can be a useful tool for proteomic analysis across various research fields.

## Conclusion

Studying protein corona formation on nanoparticles is crucial for improving human health and the clinical translation of nanomedicines, as protein coronas can directly influence how nanoparticles behave in biological systems, thereby affecting their safety, efficacy, and targeting capabilities. Despite the protein corona being a critical factor determining the properties of nanoparticles in biological environments, current protein corona data analysis relies on fragmented workflows or is limited to simple descriptive reporting, limiting the ability to derive meaningful functional and experimental interpretations from the data. Additionally, proteomics analysis has been a time-consuming process, requiring researchers to jump between separate databases and handle data plotting individually. ProCAST solves this by streamlining proteomic and sample metadata integration, database incorporation, analysis, and multi-level visualizations into a single workflow. This allows wet-lab and non-bioinformatics researchers to skip the tedious manual steps and save valuable time, helping them move faster to the next experimental phase. While universal standardization in protein corona research remains a challenge across laboratories, ProCAST provides a more consistent, less user-biased analysis framework at the data interpretation stage. Since the way we process and visualize data can yield different biological and experimental insights, we expect ProCAST to impact the nanomedicine community by enabling the rational design of drug delivery systems. Future work should include broader benchmarking across diverse nanoparticle and biological fluid systems, validating hypothesis-generating observations through follow-up experiments, and further developing the graphical user interface to make the platform more widely accessible.

## Acknowledgements

S.D.G. acknowledges financial support from the Burroughs Wellcome Fund Postdoctoral Enrichment Program (Request ID # 1021694). C.A.K. acknowledges financial support from the National Institute of General Medical Sciences (R35GM147164). Figure 1 made using BioRender. The authors would like to acknowledge N. Daniel Hills for initial organization/coding and coming up with the name “ProCAST”.

## Author Contributions

H.M.: coding, data generation, data analysis, original manuscript draft, manuscript revision and editing M.L.: original app development and coding

A.K.: original manuscript draft (introduction), dataset scrubbing

C.A.K.: coding, data generation, data analysis, initial manuscript draft, manuscript revision and editing, funding acquisition, supervision

S.D.G: inception of “ProCAST” idea, data analysis, data interpretation, initial manuscript draft, manuscript revision and editing, funding acquisition, supervision

